# Remodeling of tRNA modification in *Trypanosoma cruzi* life forms

**DOI:** 10.1101/2025.06.07.658256

**Authors:** Herbert G. S. Silva, Janaina de F. Nascimento, Marilene S. Braga, Ariel M. Silber, Matthew K. Waldor, Julia P. C da Cunha, Satoshi Kimura

## Abstract

*Trypanosoma cruzi,* the etiological agent of Chagas disease, infects millions of people in the Americas. This parasite undergoes drastic changes in its morphology and metabolism between infective and noninfective forms through global remodeling of its proteome. Chemical modification of tRNA (tRNA modification) contributes to the control of protein expression by modulating the codon decoding process. However, knowledge of tRNA modification profiles, the enzymes that create modifications and their regulation in different cellular conditions is largely restricted to relatively few model organisms. Here, we profile tRNA modifications in both infective and noninfective forms of *T. cruzi* to probe their dynamic changes. Genome mining of tRNA modifying enzymes identified 66 putative tRNA modifying enzymes in *T. cruzi*, each responsible for at least one of fifty-seven modifications. tRNA sequencing detected reverse transcription-derived signatures at 182 sites in *T. cruzi* tRNAs that are likely derived from 18 tRNA modifications. tRNA modifications and tRNA modification enzymes are differentially modulated across the life stages of *T. cruzi.* We found that hydroxywybutosine (OHyW) at position 37 on tRNA^Phe^ had a reduced level in the infective form (metacyclic trypomastigote) and the associated modification enzyme Tyw1a exhibited reduced expression in this stage. Knockout of Tyw1a increased the differentiation from epimastigote (noninfective form) to metacyclic trypomastigote, suggesting that OHyW37 modification levels control the rate of metacyclogenesis. Overall, our findings suggest that the global regulation of tRNA modifications in the life stages of *T. cruzi* plays a critical role in the differentiation of this parasite.

## Introduction

The Trypanosomatidae is a family of protozoa that includes human pathogens such as *Trypanosoma brucei*, *Leishmania major*, and *Trypanosoma cruzi*. *T. cruzi*, the causative agent of Chagas disease, poses a significant global health concern, impacting approximately seven million people worldwide ^1^. The process of differentiation between *T. cruzi* life forms involves alterations in their morphologies, metabolic patterns, and gene and protein expression ^2–7^.

*T. cruzi* alters global gene expression profiles to survive and adapt under different conditions, such as variations in pH, osmolarity, temperature, and high levels of oxidative stress ^8–10^. Since this parasite relies on polycistronic transcription where control of expression of individual genes is limited, the regulation of protein expression is predominantly governed by posttranscriptional mechanisms ^11^. Recently, we suggested that this parasite may harbor a new layer of regulatory control of protein expression based on tRNA availability and anticodon::codon pairing modes. In this system, highly expressed mRNAs show greater coadaptation to tRNA abundance and preferentially use codons that favor Watson-Crick pairing. In contrast, less expressed mRNAs exhibit lower coadaptation to tRNA abundance and tend to use codons that rely more on wobble pairing with tRNAs. ^12^.

tRNA modifications have become increasingly recognized as an important layer of regulation for protein synthesis. Through modulating the decoding ability and stability of tRNAs, tRNA modifications play a key role in the survival of organisms ^13–17^. Several studies have linked the dynamic changes in tRNA modification to critical biological processes, such as cancer progression, bacterial virulence, and the ability of organisms to adapt in extreme or hostile environments ^18–21^. The impact of tRNA modification on the efficiency of translating specific classes of proteins depends on the type and position of specific modifications. Therefore, profiling types and positions of tRNA modifications is essential to understand their roles in translation modulation. tRNA modifications are present across all kingdoms of life but vary among organisms ^22^. However, the profiles and functions of tRNA modifications are still largely undefined in many organisms.

Among Trypanosomatids, some tRNA modifications have been well studied in *T. brucei*, where they are reported to regulate the subcellular localization of tRNAs, codon recoding, and mitochondrial translation ^23–29^ However, in both *T. brucei* and *T. cruzi*, the full repertoire of tRNA modifications and their global changes across different life stages have not been evaluated.

tRNA modification sites can be predicted and identified by multiple methods ^30^. *In silico* search for homologs of known tRNA-modifying enzymes allows prediction of modifications based on known activities of tRNA-modifying enzymes in other organisms ^31^. However, this method cannot predict the modifications that are not observed in other organisms or sites of modifications that can vary among different organisms due to the variations in the substrate specificity of the homologs of tRNA modifying enzymes. tRNA sequencing (tRNA-seq) enables unbiased rapid prediction of tRNA modification sites in all tRNA species, based on the detection of reverse transcription (RT)-derived signatures ^32^. Some modifications on tRNAs inhibit Watson-Crick base pairing, which increases the frequency of misincorporation (MI) of incorrect bases and premature termination (TE) during the cDNA synthesis process ^32^. The change of tRNA modification at the specific site can be detected by the changes in these RT-derived signatures.

Here, coupling tRNA-seq and bioinformatic searches for tRNA-modifying enzyme, we profiled tRNA modifications in *T. cruzi*. tRNA modifications conserved in other eukaryotes as well as Trypanosoma specific signals were uncovered by tRNA-seq. Comparison of RT-derived signatures in tRNA-seq data from infective (metacyclic trypomastigote – MT, and tissue cultured trypomastigote - TCT) and noninfective (epimastigote-EPI) forms revealed that tRNA modification profiles are dynamic across *T. cruzi* differentiation. A subset of the dynamic changes of tRNA modification frequencies were correlated with changes in the expression levels of tRNA modifying enzymes. Finally, we found that the deletion of the tRNA modifying enzyme Tyw1a, which is associated with the OHyW37 modification pathway on tRNA^Phe^, increases parasite differentiation.

## Materials and methods

### Cell culture

The EPI forms of *T. cruzi* (Dm28c or CL Brener strains) were cultured at 28°C in Liver Infusion Tryptose (LIT) medium supplemented with 10% fetal bovine serum (FBS - Vitrocell), 0.4% glucose, 0.1 µM hemin, and 60 mg/mL penicillin G, following the protocol by Camargo et al., 1964 ^33^. The MT forms were obtained using the method described by Contreras et al., 1985 ^34^, with some modifications. Briefly, EPIs in the exponential growth phase (4x10 parasites/mL) were cultured for four days until reaching the stationary phase (5-6x10 parasites/mL). The parasites were then resuspended at a concentration of 5x10 parasites/mL in Triatomine Artificial Urine (TAU) medium (190 mM NaCl, 17 mM KCl, 2 mM CaCl, 2 mM MgCl, and 8 mM phosphate buffer, pH 6.0) and incubated for 2 hours at 28°C. Afterwards, the parasites were diluted to 5x10 /mL in TAU 3AAG medium (containing 10 mM L-proline, 50 mM L-glutamate, 2 mM L-aspartate, and 10 mM glucose) and kept at 28°C in a CO incubator. The MTs were collected from the culture supernatant after 76 hours of incubation and purified using DEAE-Cellulose resin (Sc-211213). The TCT forms of the Dm28c strains were collected from the supernatant of LLCMK2 cells one-week post-infection (at a ratio of 1:40, cells to parasites), following the method described by Nogueira et al, 1976 ^35^. The LLCMK2 cells were cultured in DMEM (Gibco) supplemented with 3.7 g/L NaHCO, 0.059 g/L penicillin G, 0.133 g/L streptomycin, and 10% fetal bovine serum (FBS) at 37°C with 5% of CO.

### tRNA sequencing, processing and analysis

tRNA-seq for TCT samples was conducted together with EPI and MT samples as previously described ^12^. Briefly, total RNA from *T. cruzi* from the Dm28c strain (in biological duplicates) was isolated using Trizol (Invitrogen), followed by the tRNAs purification and sequencing based on the method of Kimura et al., 2020 ^32^, with slight modifications described at Silva et al., 2024 ^12^. The tRNA samples were sequenced on an Illumina NextSeq 1000 system (single-end). FASTQ files from EPI and MT were obtained from PRJNA112437 and re-reprocessed together with FASTQ files from TCT. The adapter sequences (AGATCGGAAG) were removed from the FASTQ files using the cutadapt v.2.8 tool ^36^, followed by quality trimming using TrimmomaticSE v.0.39 (Bolger et al., 2014) with HEACROP:2 settings. The reads were then mapped using bowtie 1.3.1 ^38^, and mpileup files were generated using the samtools mpileup command (options: -A –ff 4 -x -B -q 0 -d 10000000 -f). The frequency of misincorporation was calculated in each mpileup file using a Python script to find the tRNAs Modifications ^32^. The coverage values of the 5’ ends of mapped reads were obtained using the bedtools genomecov command (option: -d −5 -ibam). To calculate the termination frequency, the number of 5’ ends at any position was divided by the total number of reads upstream^32^. The frequencies of misincorporation and termination were visualized using R (v.4.3.1) in a heatmap.

### Mass spectrometry

Five hundred ng of tRNA fraction purified from the total RNA extracted from *E. coli* grown in LB medium and *T. cruzi* in the exponential phase, was digested by 0.05 unit of Nuclease P1 (US Biological) and 0.1 unit of Phosphodiesterase I (Sigma) in 20 µl aliquot containing 50 mM NH_4_OAc pH 5.3, and 1 mM ZnCl_2_, at 37°C for 1 hr, followed by the addition of 2 µl of 1 M Tris-HCl pH 8.0 and 1 µl of 1 U/µl phosphatase (Sigma) for dephosphorylation at 37 °C for 30 min. After removal of the enzymes by using a 10K filter column, 5 µl of nuclease digest was loaded to LC-MS. UltiMate3000RSLCnano ultra-high-performance liquid chromatography (uHPLC) system (Thermo Scientific) bearing Synergi Fusion-RP column (100×2 mm, 2.5 µm, Phenomenex) at 40 °C with a flow rate 0.2 ml/min with a solvent system consisting of 5 mM NH_4_OAc (Buffer A) and 100% Acetonitrile (Buffer B). The gradient of acetonitrile was as follows: 0-2 min: 0.5% B, 12-15 min: 95% B, 17-20 min: 0.5% B. The eluent was ionized by an electrospray ionization source and injected into a ZenoTOF 7600 (SCIEX) with the positive mode of MRMhr scan. The voltages and source gas parameters were as follows: spray voltage; 5500 V, source name; TurbolonSpray, curtain gas; 35, CAD gas; 7, ion source gas 1; 40 psi, ion source gas 2; 50 psi, ion source temperature; 450 °C. OHyW and m^5^U signals were detected by the following mass transition; OHyW: *m/z* 525.194511393.1523 and m^5^U: *m/z* 259.0930 11127.0508.

### Identifying tRNA-modifying enzymes in Trypanosoma

A FASTA file containing the amino acid sequences of 216 enzymes involved in tRNA modifications in prokaryotes and eukaryotes was obtained from MODOMICS ^39^ except for cysteine desulfurase from *T. brucei* (Tb927.11.1670) ^40^ and a DTW domain containing protein (DTWD2 -Tb927.3.4690) ^41^. This file includes 216 of known and putative tRNA modifying enzymes present in different organisms. Subsequently, the Local Alignment Search Tool (BLAST) was applied to the genome of the Dm28c strain version 2018 (Available at https://tritrypdb.org/tritrypdb/app/workspace/blast/new) to identify the homologous genes in *T. cruzi*. Alignments with an E-value ≤1E-10 were considered to be homologous genes in *T. cruzi.* The tModBase database ^42^ was used as an auxiliary tool to identify the modification sites on tRNA for the enzymes characterized in this study. Orthologous tRNA-modifying enzymes in *T. brucei* TREU927 were identified using the ‘strategies’ and “orthology phylogenetic profiles” tools available in TriTrypDB (https://tritrypdb.org/tritrypdb/app). The cellular localization of enzymes associated with tRNA modifications and their impact on survival in *T. brucei* was examined using the platform (http://tryptag.org/) ^43^and by consulting the list of proteins provided by Asford et al., 2011 ^44^, respectively.

### Generation of null mutants using CRISPR-Cas9 and addback cell lines

Parasites CL Brener (EPI) expressing the Cas9 protein endogenously were used for genome edition ^45^. Primers listed in Supplementary Table 3 were designed using the EuPaDGT tool ^46^ to obtain the sgRNA sequences for targets: Tyw1a: TcCLB.508277.60 and TcCLB.503543.4. The sgRNAs were amplified by PCR using Platinum Taq DNA Polymerase (Invitrogen). For amplifying the DNA donor, 30 bp homologous sequences located up and downstream of sgRNA binding sites of the genes of interest were added in the template primers to replace the genes of interest with the blasticidin or puromycin resistance genes. Primers were designed manually (in Supplementary Table 3) and the plasmids pJET-blast or pT-puro ^45^ were used as templates for the PCR using 3 U Platinum Taq DNA Polymerase High Fidelity (Invitrogen) in a total reaction volume of 120 μL. To generate addback cell lines, the coding sequences of Tyw1a were amplified by PCR using specific primers flanked by restriction sites (Supplementary Table 3). The Tyw1a PCR product was gel purified using QIAquick Gel Extraction Kit according to manufacturer’s instructions, digested with EcoRI Fast digest and XhoI Fast digest (Thermo Fisher Scientific) and ligated with similarly digested pTEX_puroR treated with FastAP Thermosensitive Alkaline Phosphatase (Thermo Fisher Scientific). Before transfection, sgRNA and DNA donor PCR products and 25 μg of the plasmids were purified, precipitated and resuspended in 5 μL of sterile ultrapure water each. To parasite transfections, 4 × 10^7^ of EPI were treated with 20 mM hydroxyurea (Sigma) for 18 hours ^47^. Then, the parasites (4 × 10^7^) were resuspended in 1 mL of transfection buffer (90 mM sodium phosphate, 5 mM potassium chloride, 0.15 mM calcium chloride, 50 mM HEPES, pH 7.2), along with the DNA to be transfected in a 0.2 cm cuvette (BioRad) using the Nucleofector 2b device (Lonza) with a single pulse using program U-033. The modified parasites were selected in the presence of antibiotics and subsequently cloned in 96-well plates. Null and addback parasites lineages were confirmed by PCR using specific pairs of primers described in Supplementary Table 3. Agarose gel electrophoresis was used to visualize the positive PCR amplifications.

### Western blot

Total protein extracts were separated by 10% polyacrylamide gel electrophoresis (SDS-PAGE) and transferred to a nitrocellulose membrane (BioRad) using the TransBlot Turbo system (BioRad), as described by ^48^ and following the manufacturer’s instructions. The membrane was blocked for 1 hour in PBS-T (0.1% PBS-Tween-20) solution containing 5% skim milk. After blocking, the membrane was incubated with a primary anti-6xHisTag monoclonal antibody (Invitrogen, MA121315) at a 1:1000 dilution in PBS-T solution containing 3% skim milk, with gentle agitation overnight at 4°C. Next, the membrane was washed three times for 5 minutes with PBS-T. Subsequently, the membrane was incubated for 1 hour with the secondary antibody conjugated to horseradish peroxidase (HRP, dilution 1:2000; GE Healthcare), followed by additional washes as described above. Finally, the assays were developed using chemiluminescence with the ECL Super Signal West Pico Chemiluminescent Substrate (Thermo Scientific) according to the manufacturer’s instructions and visualized using the ChemiDoc Imaging System transilluminator (BioRad).

### RT-qPCR

Total RNA extractions from *T. cruzi* null and addback lineages were performed using Trizol (Invitrogen) according to the manufacturer’s instructions. The reverse transcription reaction was performed using RNA previously treated with DNAse I, Amplification Grade (Invitrogen) and the SuperScript™ III First-Strand Synthesis (Invitrogen™) using 50 ng Oligo(dT)12-18 (Invitrogen), following the manufacturer’s instructions. For each sample, a control without reverse transcriptase (-RT) was prepared. qPCR reactions were performed using the 5x HOT FIREPol® Probe Universal qPCR Mix (SolisBioDyne) including 4 µL of reagent mix, 400 nM of primers, 1x SYBR Green, 1 µL of cDNA, and nuclease-free water. All samples were analyzed in technical triplicate, with parallel reactions performed without cDNA or with -RT samples as negative controls. Amplification conditions were carried out on the StepOnePlusTM Real-Time PCR system (Applied Biosystems®) as follows: initial denaturation at 95°C for 10 minutes followed by 40 cycles of denaturation at 94 °C for 15 seconds and annealing at 60 °C for 1 minute, and a final melting curve from 60 °C to 95 °C. Primers were designed using Primer3 (https://primer3.ut.ee/) ^49^ the sequences of Tyw1a, and GAPDH (TcCLB.506943.50) as templates (Supplementary Table 3). The gene glyceraldehyde 3-phosphate dehydrogenase (TcGAPDH) was used as a normalizer for calculating relative expression. Data were analyzed using the StepOne Software v2.3. Variations in transcript expression were determined using the 2(-ΔΔCt) method ^50^.

### *In vitro* metacyclogenesis for *T. cruzi* null and addback strains

To conduct *in vitro* metacyclogenesis assays, two different methods were used. In the first method ^51^, EPIs in the exponential growth phase were used to initiate a culture with an initial concentration of 5 x 10^6^ parasites ml^-1^, which was maintained in LIT as previously described for 6 days. After, the parasites were washed with PBS 1x and then transferred to TAU medium (190 mM NaCl, 17 mM KCl, 2 mM MgCl_2_, 2 mM CaCl_2_, 8 mM potassium phosphate buffer, pH 6) at a concentration of 5 x 10^7^ parasites ml^-1^ and maintained for two hours at 28°C. The parasites were subsequently transferred to TAU 3AAG (TAU supplemented with 10 mM glucose, 2 mM aspartic acid, 50 mM glutamic acid, 10 mM proline) and maintained for 8 days in a CO_2_ incubator at 28°C. The presence of MT was checked daily by counting in a Neubauer chamber. In the second protocol ^52^, 1 mL of EPI culture in the exponential growth phase (5 x 10^7^ cell ml^-1^) was transferred to a culture flask containing 10 mL of RPMI (Vitrocell; prepared according to the manufacturer’s instructions, without the addition of FCS and pH adjustment). The flasks were kept undisturbed at a 30° angle in a CO_2_ incubator at 28°C for 8 days. After 8 days, the number of MT in the supernatant was counted using a Neubauer chamber.

### Graphs and statistical analysis

All graphs and statistical analysis were performed using GraphPad Prism v10.

## Results

### *In silico* identification of tRNA modifying enzymes in Trypanosoma and predicting their essentiality and localization

We compiled a list of 216 protein sequences of known and putative tRNA-modifying enzymes present in various eukaryotic and prokaryotic species, ranging from human to *Escherichia coli* (Supplementary Table 1A). BLAST was employed to search for homologs of these tRNA modifying enzymes in the *T. cruzi* genome. Using an E-value of 1e-10 as a cutoff, we identified sixty-six homologs of tRNA-modifying enzymes in *T. cruzi*; these enzymes mediate various tRNA modifications, including dihydrouridinylation, thiolation, aminocarboxypropylation, methylation and acetylation (Figure 1A and Supplementary Table 1B). Some tRNA modifying enzymes have multiple paralogs in *T. cruzi*. For example, the methyltransferase Ab140 from yeast (Gene ID= Q08641) has two orthologs in *T. cruzi*, Ab140a (C4B63_2g720) and Ab140b (C4B63_58g34) (Figure 1A and Supplementary Table 1B). We added suffixes (a, b, or c) to the end of the enzyme name to distinguish the candidate paralogs identified in this study. All these tRNA modifying enzymes have homologs in *T. brucei* (Supplementary Table 1C), which displays 94% similarity and 57% identity with the *T. cruzi* genome ^53,54^, suggesting that tRNA modification profiles are conserved in different species of Trypanosoma.

**Figure 1.**
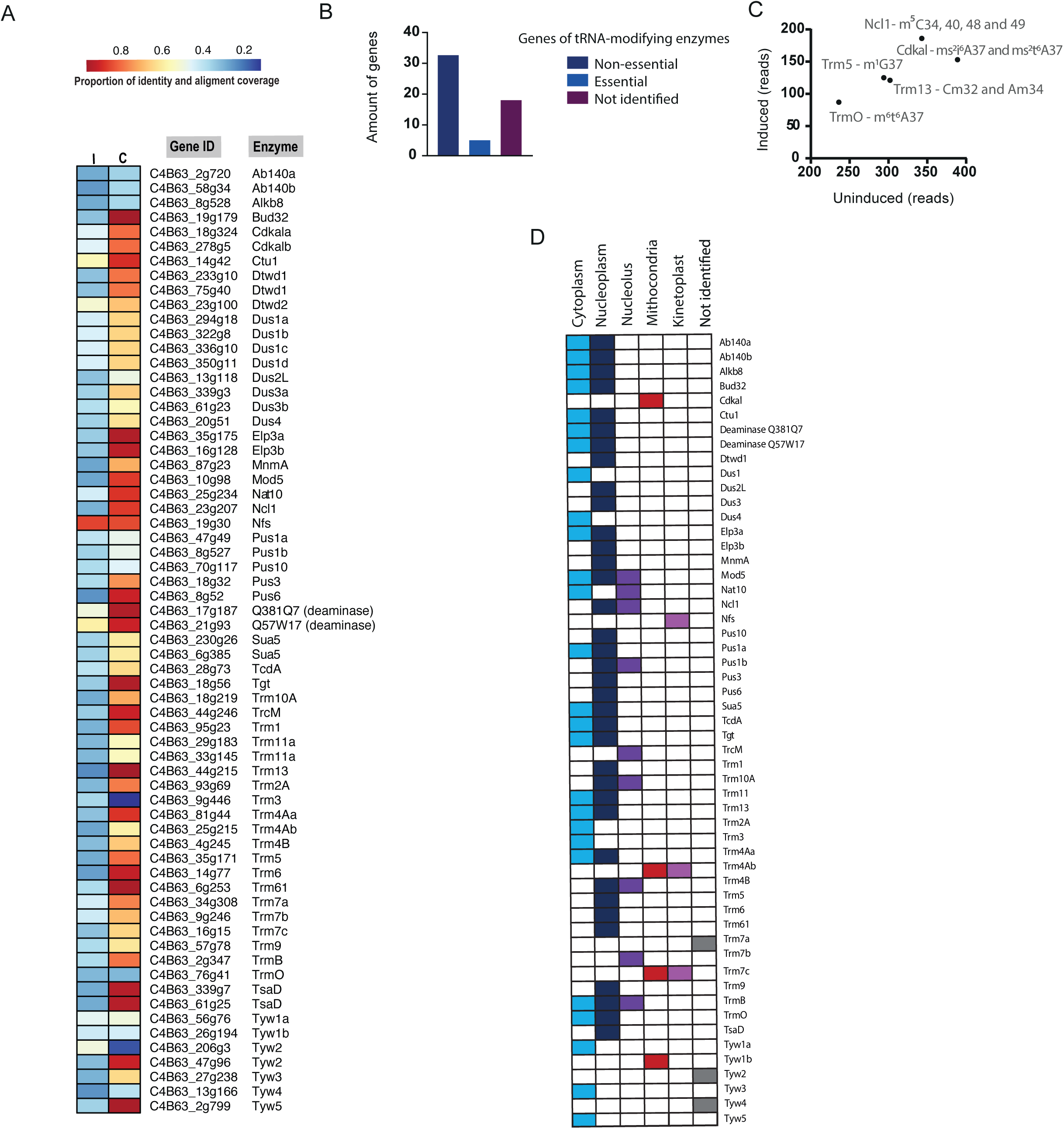
Identification, essentiality, and localization of tRNA-modifying enzymes in Trypanosoma. **A)** BLAST search of tRNA modifying enzymes in *T. cruzi*. Proportion of alignment coverage (C) and identity (I) between subject protein sequences from the *T. cruzi* genome (Dm28c – version 2018) and known tRNA-modifying enzymes listed in Supplementary table 1B. The heatmap displays the gene ID and enzyme names of the candidate homologs in *T. cruzi* identified. E-values ≤1e-10 were used as a threshold for candidate selection. **B**) Essentiality of tRNA-modifying enzymes in Trypanosoma. The number of tRNA-modifying enzyme genes identified in public RNA Interference Target Sequencing (RIT-seq) data ^44^ associated with *T. brucei* survival during the differentiation of procyclic to bloodstream trypomastigote forms. **C)** Read counts normalized to Reads Per Five Million (RPFM), representing RNAi-induced insertions targeting tRNA-modifying enzyme mRNAs that influence *T. brucei* survival during differentiation. Read counts in induced (y-axis, silenced) and non-induced (x-axis, not-silenced) parasites were analyzed using the RIT-seq technique. **D**) **S**ubcellular localization of tRNA-modifying enzymes in the *T. brucei*: cytoplasm, nucleoplasm, nucleolus, mitochondria, and kinetoplast. The cellular localization of some proteins, for which no public data is available, is shown as ‘not identified’.

To gain further insights into the impact of tRNA-modifying enzymes on *Trypanosoma* fitness, we analyzed high-throughput phenotyping data, RNA Interference Target Sequencing (RIT-seq) data, available in *T. brucei*^44^. Among the fifty-six tRNA-modifying enzymes identified in *T. brucei*, no data is available for nineteen (34%) (Figure 1B and Supplementary Table 1D). Of the enzymes with RIT-seq data, thirty-two (86%) were classified as non-essential, while five (14%) were considered essential during the *T. brucei* life stages, such as Trm13 and TrmO, involved with m^1^G37 and m^6^t^6^A37modifications, respectively (Figure 1C). These results indicate that specific tRNA modifications play a key role in *Trypanosoma* fitness.

tRNA modifying enzymes are known to be found at various subcellular locations. To evaluate their cellular localization, we interrogated the *T. brucei* genome-wide database of subcellular protein localization ^43^. We found that most of the tRNA-modifying enzymes are in nucleoplasm and cytoplasm (Figure 1D), whereas some enzymes are localized in organelles. Trm7c and Trm4Ab associated with Gm and m^5^U, respectively, are predicted to localize in both mitochondria and kinetoplast (Figure 1D). Curiously, we identified some paralogs, such as Trm4Aa and Trm4Ab, which exhibit distinct subcellular localization patterns, where Trm4Aa can be found in cytoplasm/nucleoplasm and Trm4Ab in mitochondria/kinetoplast. These data suggest that these homologous enzymes catalyze the same modification in tRNAs, but in different cellular compartments.

### Profiling tRNA modification sites in *T. cruzi*

To profile tRNA modification sites in *T. cruzi*, we employed tRNA-seq. tRNA from exponentially growing non-infective state (EPI form) was purified and sequenced and exhibited high sequence quality and read mapping across replicates (Supplementary Figure 1A-B). Several tRNA modifications perturb reverse transcription during cDNA synthesis, which leads to the misincorporation of incorrect bases and premature termination of reverse transcription, which we call reverse transcription-derived signatures (RT-derived signatures). Therefore, RT-derived signatures detected in tRNA-seq data likely correspond to modified sites as previously published in prokaryotes ^32,55^. The misincorporation and termination signals in *T. cruzi* tRNAs are shown in Figure 2A and Supplementary Figure 2.

**Figure 2.**
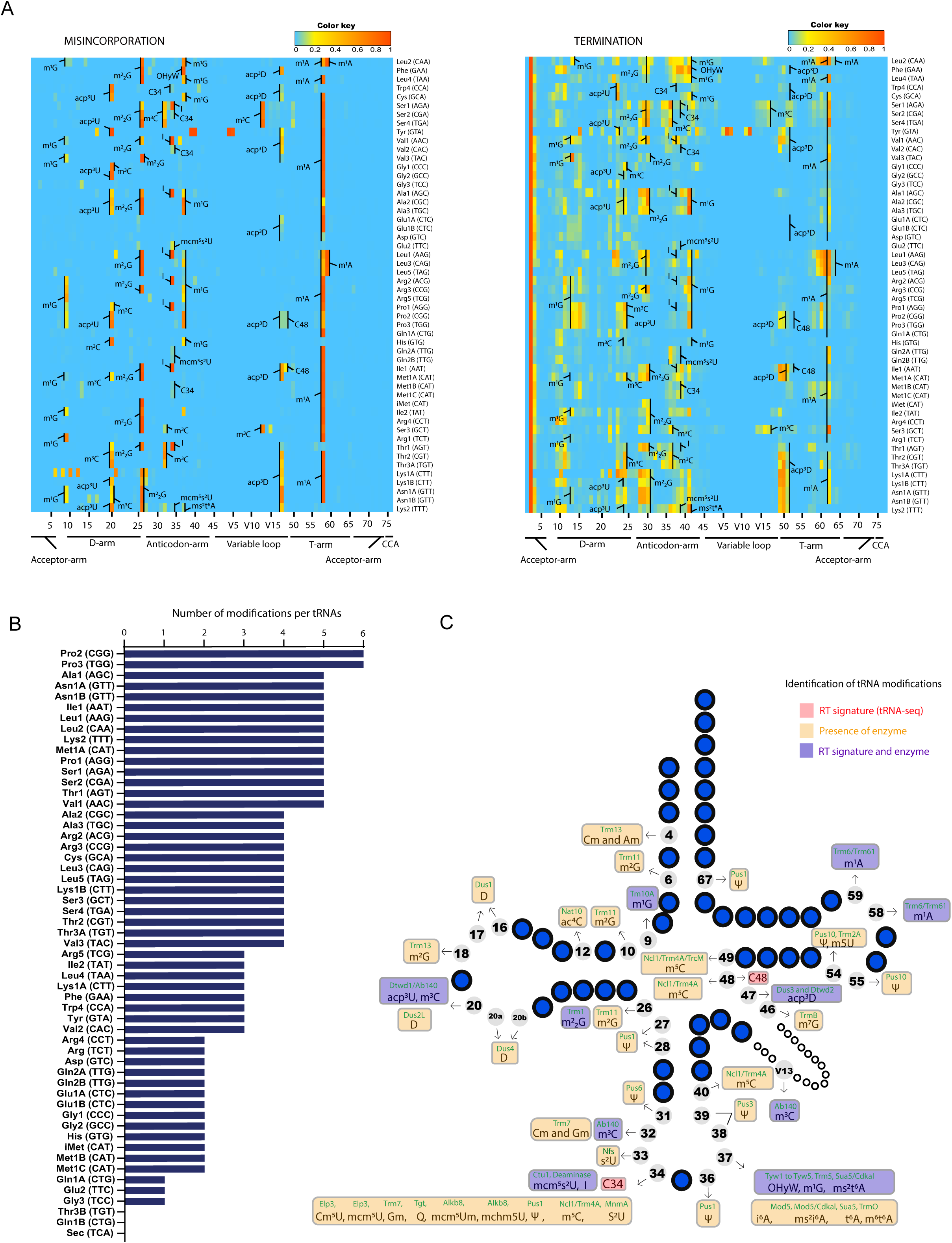
The tRNA modification profiles in *T. cruzi* EPI life form. **A)** Heatmaps display misincorporation and termination frequencies calculated from tRNA-seq results from the *T. cruzi* EPI form. Predicted tRNA modifications on RT-derived signatures are shown. The position and species of tRNA are indicated in the x- and y-axis, respectively. **B)** Number of modifications identified in different tRNA species using tRNA-seq data. **C)** Overview of all tRNA modifications detected by tRNA-seq and predicted based on the presence of tRNA-modifying enzymes. The tRNA-modifying enzymes are highlighted in green above their respective modifications. Modifications predicted by both RT-derived signature and the presence of the corresponding enzyme are shown in purple.

Many RT-derived signatures are consistent with conserved modification sites in eukaryotes, such as human and yeast ^42^. The sites and modifications observed in the *T. cruzi* profiles include m^1^G at position 9, m^3^C and acp^3^U at position 20, m^2^_2_G at position 26, m^3^C at position 32, 20 and V13, I at position 34, m^1^G, OHyW, and ms^2^t^6^A at position 37, and m^1^A at position 58 and 59 (Figure 2A). Furthermore, the homologs of tRNA modifying enzymes that synthesize these modifications were also identified in *T. cruzi*, such as m^1^G9: TRMT10A; acp^3^U20: DTWD1; m^2^_2_G26: TRMT1; m^3^C32: ABP140; I34: ADAT2 – deaminase; m^1^G37: TRMT5; OHyW37: Tyw1,2,3,4,5; ms^2^t^6^A37: CDKAL, SUA5; m^3^C20, m^3^C32, V13: ABP140 (Figure 1A, Figure 2C and Supplementary Table 1B). These observations support the assignment of tRNA modifications to *T. cruzi* RT-derived signatures.

We detected misincorporation and termination signatures that are derived from the modifications observed in *T. brucei*, at position 47 in twenty-two tRNA species, such as tRNA^Lys1A(CTT)^, tRNA^Asn1B^ ^(GTT)^, tRNA^Lys2^ ^(TTT)^ (Figure 2A and Figure 2C). In *T. brucei*, tRNA^Lys(TTT)^ is modified at position 47 by a unique modification acp^3^D ^29^, suggesting that the RT-derived signature detected in *T. cruzi* is likely derived from acp^3^D.

Some RT-derived signatures in *T. cruzi* are likely derived from the modifications in mitochondrial tRNAs. Unlike mammals and yeast, Trypanosoma utilizes nuclear-encoded tRNAs not only in the cytoplasm but also in mitochondria ^56,57^. A fraction of tRNAs is imported into mitochondria and some of them are further modified. Thus, some of the RT-derived signatures are likely derived from the fraction of mitochondrial tRNAs. In *T. brucei*, tRNA^Trp4(CCA)^ is imported into mitochondria, and its wobble position is edited from C-to-U leading to decoding the termination codon (UGA) as tryptophan ^58^. The misincorporation signals at position 34 observed in *T. cruzi* tRNA^Trp4(CCA)^ (Figure 2A) suggest that *T. cruzi* tRNA^Trp4(CCA)^ is also edited at position 34 for UGA codon recoding. Additionally, the wobble position of tRNA^Met^ is modified into formyl cytidine (f^5^C) in mitochondria in mammals to decode two methionine codons, AUG and AUA, in mitochondria ^59^. Misincorporation signals at position 34 in tRNA^Met1B(CAT)^, tRNA^Met1C(CAT)^ (Figure 2A), and the presence of a homolog of ALKBH1 (C4B63_7g420), which synthesizes f^5^C, in *T. cruzi* suggests that f^5^C is utilized to expand the codon decoding ability of tRNA^Met^ in Trypanosoma mitochondria.

In addition, we observed some RT-derived signatures in tRNAs from *T. cruzi* that are not observed in other organisms, such as misincorporation at position 34 in tRNA^Ser2(CGA)^ and tRNA^Val2(CAC)^, and at C48 in tRNA^Pro2(CGG)^, tRNA^Pro3(TGG)^, and tRNA^Ile1(AAT)^. These signatures likely represent modifications exclusively present in *T. cruzi* and other trypanosomatids.

While we found *T. cruzi* homologs of tRNA-modifying enzymes *in silico* in *T. cruzi*, some expected tRNA modifications were undetectable by tRNA-seq due to the absence of RT-derived signatures. We found 42 enzymes that likely synthesize tRNA modifications in *T. cruzi,* but their modification sites were not detected by tRNA-seq, as they do not inhibit Watson-Crick base pairing, such as D20, m^5^U54, Q34 and cm^5^U34 (Figure 2C, Supplementary Table 1B and Supplementary Table 2B).

Finally, we found distinct number of modifications per tRNA species. For instance, tRNAGly^3(TCC)^ and tRNA ^Pro2(CGG)^ have one and six modifications, respectively, detected by tRNA-seq (Figure 2B). The m^1^A58 is the most predominant modification found in forty-six tRNA species, while OHyW37 and ms^2^t^6^A37 are present in a single tRNA species (Supplementary Figure 3).

### tRNA modification levels change in different *T. cruzi* life forms

tRNA modification levels can change under different environmental conditions for optimizing the synthesis of specific proteins ^32,60^. *T. cruzi* changes their life forms in the face various environmental pressures; the different forms include a noninfective replicative EPI form, and infective and cell-cycle arrested TCT and MT forms. ^7–10,61^. We hypothesized that differential expression of tRNA modifying enzymes could contribute to the optimization of protein expression profiles in different *T. cruzi* life forms via alteration of tRNA modification profiles. To test this idea, we investigated whether tRNA-modifying enzymes are differently expressed in their replicative (EPI) and non-replicative (MT) forms using publicly available Ribo-seq data ^12^.

Among the sixty-five *T. cruzi* tRNA-modifying enzymes, seven (10%) were differentially expressed between EPI and MT forms (Figure 3A and Supplementary Table 1E). In the non-replicative MT form, all seven of these transcripts were downregulated by more than 2-fold (Figure 3A). Interestingly, we observed different trends in the expression of multiple paralogs of tRNA-modifying enzymes. The Ab140a and Ab140b, which are predicted to be localized at both cytoplasm and nucleoplasm are differently regulated in the EPI and MT forms (Figure 3A). Ab140a shows similar expression levels in EPI and MT cells, while Ab140b exhibits a drastic ∼25-fold reduction in expression in MT cells, suggesting that the frequency of m^3^C synthesized by Ab140b specifically decreased in MT form.

**Figure 3.**
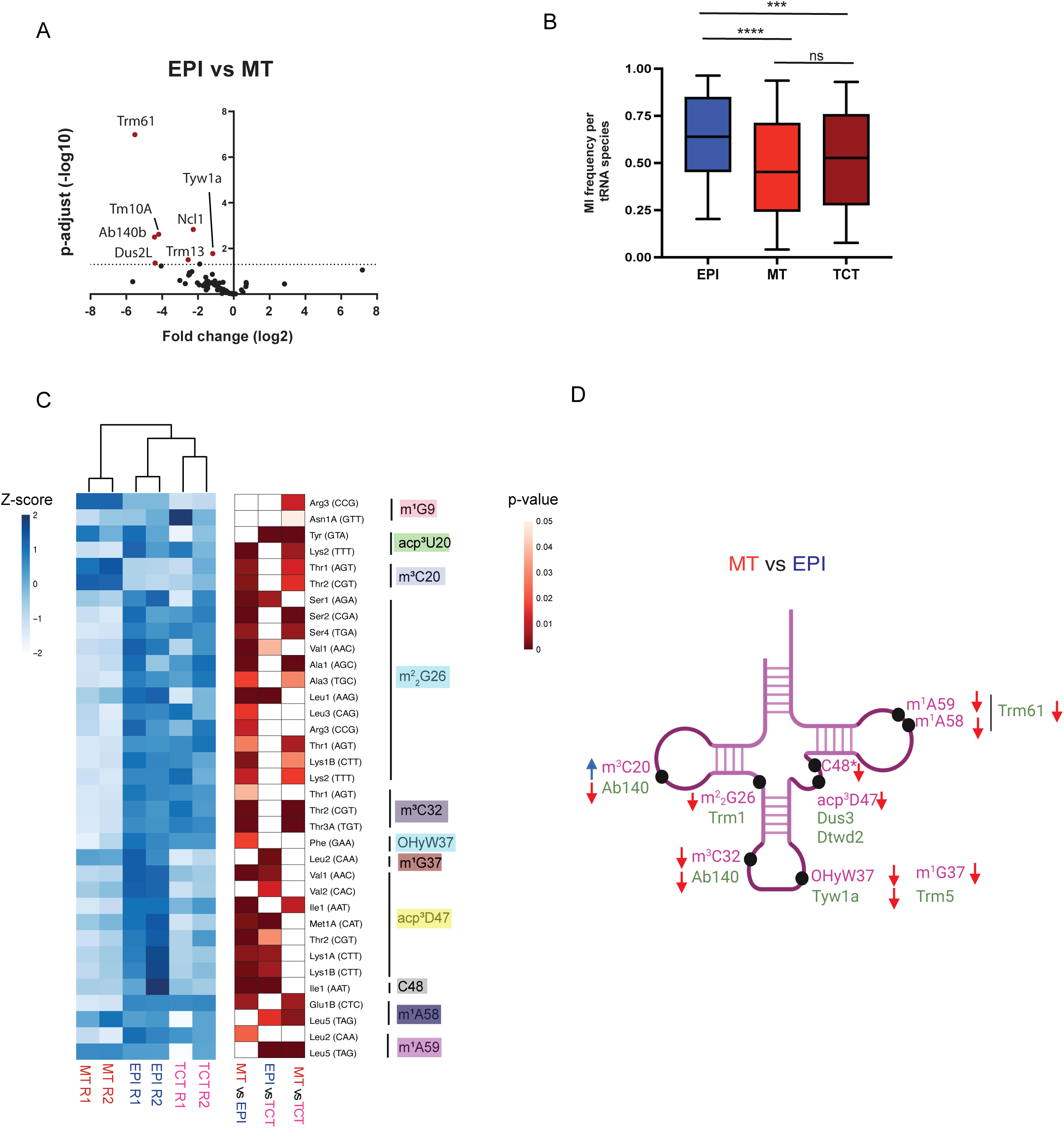
Changes in the expression levels of tRNA-modifying enzymes and the frequencies of tRNA modification across different stages of *T. cruzi*. **A)** Volcano plot showing the fold change (log_2_) and p-adjust (log_10_) of the expression levels of tRNA-modifying enzymes in EPI compared to MT using Ribo-seq data ^7^. **B)** Global changes of misincorporation rates in tRNA-seq data derived from *T. cruzi* life forms (EPI, MT and TCT). Statistical significance tests (p-value ≤ 0.05) were performed with the one-way ANOVA test. **C)** Differential tRNA modification frequencies among EPI, MT, and TCT forms. The heatmaps illustrate the z-scores of misincorporation frequencies (left) and p-values from the indicated comparisons of the misincorporation frequencies derived from tRNA modifications (right). A two-way ANOVA test was employed to identify significantly differentially tRNA modifications containing a fold change of >= 20% between life forms. **D)** Illustration containing the abundance of tRNA modification and their corresponding modifying enzymes (Trm61, Dus3, Dtwd2, Trm5, Tyw1a, Ab140a, Ab140b, Trm1) in *T. cruzi*’s life forms. The tRNA modifying enzymes are shown in green, positioned below their respective catalyzed modifications. The red arrow indicates downregulation of tRNA modifying enzymes or tRNA modification in MT compared to EPI, while blue arrow indicates upregulation in MT. Enzymes without arrows have similar abundance in both EPI and MT.

Biosynthesis of wyosine (imG) and its derivatives, including wybutosine (yW) and hydroxywybutosine (OHyW) occurs in the cytoplasm of most eukaryotes. In contrast, in *T. brucei*, two independent biosynthesis pathways for wyosine derivatives are found in the cytoplasm (OHyW) and mitochondria (imG) ^62^. Like *T. brucei*, *T. cruzi* encodes two paralogous Tyw1 loci, Tyw1a and Tyw1b, which are predicted to localize in the cytoplasm and mitochondria respectively (Figure 1D), suggesting that *T. cruzi* also possesses two parallel biosynthesis pathways for wyosine derivatives in these two cellular compartments. The level of cytoplasmic Tyw1a was 2.3-fold greater in EPI vs MT cells, while mitochondrial Tyw1b had similar expression levels in both life forms. Since these enzymes are presumably involved in the formation of wyosine derivatives, such as wybutosine and hydroxywybutosine at 37 in tRNA^Phe^, the differential expression of Tyw1a suggests that OHyW levels in cytoplasmic tRNA are regulated in a stage-specific fashion.

Next, to track the changes in the frequency of tRNA modification in different life forms of *T. cruzi*, we conducted tRNA-seq in EPI, MT and TCT forms (Supplementary Figure 1). Using misincorporation frequency as a read-out of the frequency of tRNA modification, we tracked the changes of sixteen tRNA modifications at 182 positions across all tRNA species (Supplementary Table 2C). Overall, changes in the frequency of tRNA modifications were observed among MT, TCT, and EPI forms (Figure 3B). The level of tRNA modifications from EPI cells was more similar to TCT than MT cells (Figure 3C). When analyzing individual sites, we observed changes across different types of tRNA modifications. The frequency of 12 modifications, including m^2^_2_G26, m^3^C32 and OHyW37, was comparable between EPI and TCT but reduced in MT in many tRNA species (Figure 3C-D and Supplementary Figure 4), whereas the frequency of some tRNA modifications was increased in MT (m^3^C20 in tRNA^Thr2(CGT)^ and tRNA^Thr1AGT^) compared to other forms (Figure 3C an Supplementary Figure 4). Interestingly, a drastic reduction (log_2_(fold change) ranges from −5 to −0.35) of acp^3^D47 was observed in several tRNAs, such as tRNA^Val1^ ^(AAC)^, tRNA^Lys1A^ ^(CTT)^, tRNA^Thr2^ ^(GGT)^, tRNA^Met1A^ ^(CAT)^ in the MT and TCT forms comparing to EPI (Figure 3C and Supplementary Figure 4). The acp^3^U47 is known to be important for thermal stability on tRNA ^63^, suggesting that the reduction of acp^3^D47 on tRNAs in both non-proliferative forms could result in reduced thermostability compared to tRNAs in the EPI forms.

The changes in tRNA modifications are likely at least partly caused by the differential expression of tRNA modifying enzymes in different life forms. Three tRNA-modifying enzymes downregulated in MT (Tyw1a, Ab140b, and Trm61) catalyze modifications (OHyW37, m^3^C32, m^1^A58, and m^1^A59) that exhibit a detectable reduction in their frequency in the MT form in tRNA-seq data (Figure 3C and Figure 3D), suggesting that the tRNA modification frequency is controlled by the regulation of the tRNA modifying enzyme expression. However, some significant reduction in tRNA modification levels was not explained by the expression level of their corresponding tRNA modifying enzymes. For example, the level of m^3^C20 increased in MT compared to EPI, while the expression levels of its modifying enzymes (Ab140a or Ab140b) do not change accordingly in the life forms, suggesting an alternative mechanism controls the m^3^C20 levels in MT.

### Knockout of Tyw1a promotes *T. cruzi* Differentiation

Next, we investigated whether the tRNA modifications and their associated tRNA-modifying enzymes that are differentially regulated across life stages influence *T. cruzi* differentiation. The tRNA-seq data showed that the misincorporation signal corresponding to the OHyW37 modification in MT decreased by approximately 1.23-fold compared to EPI, while the level of the Tyw1a enzyme in MT decreased by approximately 7-fold relative to EPI. Tyw1a, also known as Tyw1L, is essential for the formation of hydroxywybutosine (OHyW37) on tRNA^Phe^ in *T. brucei* ^62^. *T. cruzi* bears the other homologs required for OHyW synthesis (Tyw2, Tyw3, Tyw4, and Tyw5). Furthermore, mass spec analysis confirmed that a signal corresponding to OHyW (*m/z* 525.1945 ⟶ 393.1523) was detected in the nucleoside pool of *T. cruzi* tRNA but not in that of *E. coli* (Figure 4A), strongly supporting that *T. cruzi* also has OHyW in tRNA^Phe(GAA)^. Here, we investigated whether the deletion of Tyw1a could affect *T. cruzi* differentiation.

**Figure 4.**
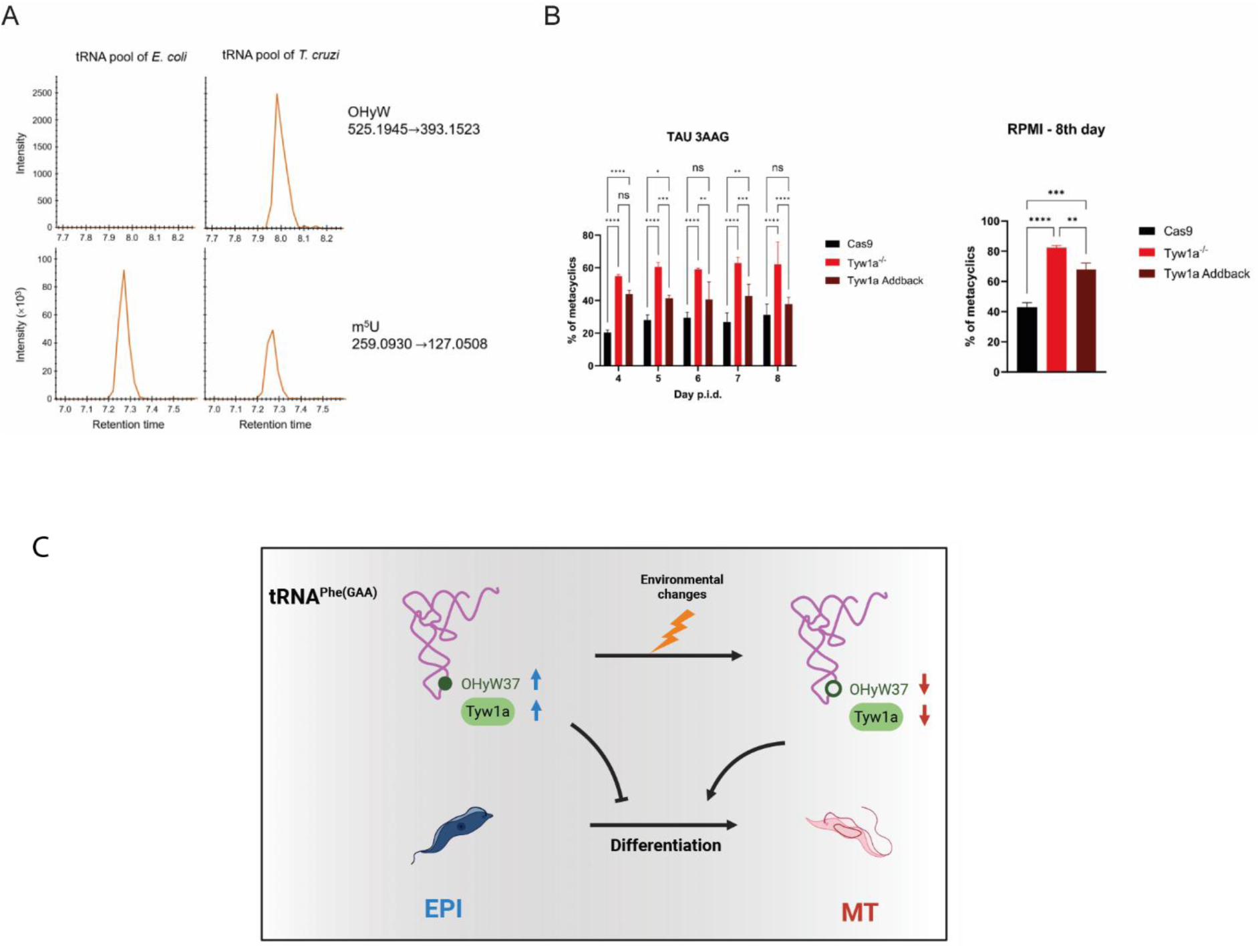
Tyw1a knockout facilitated *T. cruzi* differentiation. **A)** LC-MS analysis detected OHyW in *T. cruzi* tRNA pool (EPI) but not in *E. coli*, whereas m^5^U was detected in both tRNA fractions from *T. cruzi* and *E. coli.* The panels show the signal intensities of ions undergoing the mass transition 525.1945 IZ 393.1523, corresponding to OHyW, and 269.0930 IZ 127.0508, corresponding to m^5^U. **B)** The frequency of metacyclogenesis induced in TAU 3AAG and RPMI medium. The parental cell line (Cas9), Tyw1a knockout (Tyw1a^-/-^), and Tyw1a add-back cell lines are shown. Experiments were performed with 5 biological replicates of each cell line. The Cas9 strain expressing only Cas9 but not guide RNA was used as a control. Statistical analysis was performed using one-way ANOVA (RPMI) or two-way ANOVA (TAU 3AAG) with multiple comparisons test (a=0.05, * p ≤ 0.05, ** p ≤ 0.01, *** p ≤ 0.001). **C)** Schematic illustration containing the abundance of Tyw1a and OHyW37 modification on tRNA^phe(GAA)^ in *T. cruzi*. The red arrow indicates downregulation of Tyw1a or OHyW37 modification in MT compared to EPI, while the blue arrow indicates upregulation in EPI. Reduced Tyw1a levels facilitate the metacyclogenesis process.

We generated EPI cell lines lacking Tyw1a using the CRISPR-Cas9 technique, along with the add-back cell lines (Supplementary Figure 5). We confirmed that the KO lineages do not express the Tyw1 enzyme, while the complemented parasite expresses Tyw1 (Supplementary Figure 5C). Subsequently, the parental strain (Cas9 strains), Tyw1 KO parasites (Tyw1^-/-^), and Tyw1 add-back parasites were induced to metacyclogenesis using two different differentiation strategies (TAU 3AAG and RPMI) ^34,64^. In both conditions, the Tyw1 KO parasites exhibited more than a two-fold increase in the frequency of differentiation from EPI to MT compared to the parental strain (Cas9), while this increase in differentiation was partially or fully blocked by the ectopic expression of Tyw1a (Figure 4B). These results strongly suggest that Tyw1a and the synthesis of OHyW slow down *T. cruzi* differentiation. Given that the Tyw1 expression level and OHyW frequency were lower in the MT form, *T. cruzi* may facilitate differentiation from the non-infective EPI to the infective MT form through the suppression of Tyw1a expression and its consequent decrease in the OHyW levels (Figure 4C).

## Discussion

The profiles and biological roles of tRNA modifications remain largely unexplored across many organisms. In this study, we profiled tRNA modifications in different *T. cruzi* life forms and observed dynamic changes in the level of many tRNA modifications. We found that OHyW37, whose level was decreased in the non-replicative MT form, likely facilitates differentiation from the replicative EPI to the non-replicative MT form, suggesting that *T. cruzi* can control tRNA modification level to regulate the rate of differentiation. Other tRNA modifications that showed differential levels among different *T. cruzi* lifeforms in our data may contribute to the control of differentiation of life forms.

tRNA-seq predicted the following tRNA modification sites: m^1^G9, acp^3^U20, m^3^C20, m^2^_2_G26, m^3^C32, mcm^5^s^2^U34, I34, OHyW37, m^1^G37, ms^2^t^6^A37, f^5^C34, C-to-U34, m^3^CV13, acp^3^D47, m^1^A58, m^1^A59, C34, and C48. BLAST search found *T. cruzi* homologs of enzymes that create these modifications, supporting the mapping of these modifications. Some types of modifications generated by homologs of tRNA modifying enzymes in *T. cruzi* were not detected through tRNA-seq since tRNA-seq cannot detect modification that do not affect Watson-Crick base pairing during reverse transcription ^32,65–68^. For example, m^5^C cannot be detected through tRNA-seq, although we found three homologs (Ncl1, Trm4Aa, and Trm4Ab) of the methyltransferase that generates m^5^C. To map such additional chemical modifications would be needed. Recently, in *T. brucei*, m^5^C was mapped through the bisulfide method ^69^, revealing that *T. brucei* has m^5^C at positions observed in other eukaryotes such as 34, 48, and 49. Since *T. cruzi* and *T. brucei* have the same set of homologs of m^5^C methyltransferases, *T. cruzi* may have the same profiles of m^5^C in tRNAs. Additionally, m^5^C was found at three more positions, C12, C50, and C60, in *T. brucei* tRNAs, further suggesting that the five C5-methyltransferases identified in *T. cruzi* (C4B63_23G207, C4B63_25G215, C4B63_81G44, C4B63_44g246 and C4B63_4g245) may be involved in catalyzing these modifications in its tRNAs.

In *T. cruzi*, we detected misincorporation signatures at cytidine positions 34 and 48 (C34 and C48) by tRNA-seq in certain tRNA species, such as tRNA^Ser2^ ^(CGA)^ and tRNA^Pro2^ ^(CGG)^, respectively, that don’t correspond to known modifications in other organisms. C48 is frequently methylated to form m^5^C in eukaryotes ^42^; however, this methylation doesn’t produce RT-derived signatures due to its lack of direct effects on Watson-Crick base pairing ^70,71^. While we cannot rule out the possibility that sequence context affects m^5^C to cause RT-derived signatures, C48 may undergo other modifications that have not yet been observed. tRNA-seq also detected misincorporation at C34 in multiple tRNA species. In *T. brucei*, C34 in other tRNA species undergoes modifications that cause RT-derived signatures, such as C-to-U editing ^58^. It is possible that these modifications and editing occur in other tRNA species in *T. cruzi*. Further experiments, including mass spectrometry analysis of purified tRNAs, are necessary to identify these modification types.

One of the Trypanosoma-specific modifications is acp^3^D ^72^. While many tRNA modifications showed a decrease in misincorporation only in the MT form, which is one of two non-proliferative forms, misincorporation signatures corresponding to acp^3^D47 showed a decrease in both non-proliferative forms, MT and TCT. acp^3^D47 is predicted to be present in 22 tRNA species, with over 23% of them showing reduced expression during the parasite’s non-proliferative stages. The 3-amino-3-carboxypropyl (acp) group is a highly conserved modification in both bacteria and eukaryotes, attached to the N3 atom of uracil (acp^3^U) at positions 20 and 47 ^39,72,73^. In *T. cruzi*, we detected RT-derived signatures for acp3 at positions 20 and 47 on tRNAs; however, we predict that at position 47 this modification occurs on a dihydrouridine (D) rather than a uridine. The D base is not detected by tRNA-seq, as it does not induce misincorporation or premature termination during cDNA synthesis ^32^. The prediction of modifications at the D47 base is based on two pieces of evidence: i. tRNAs with acp^3^ at position 47 within a D base instead of a U was detected in tRNA^Lys(TTT)^ from *T. brucei* ^29^; ii. the presence of the homologous DUS3L, associated with the U47-to-D47 modification ^74^ along with the enzyme DTWD2 involved in acp3 ^75^ in *T. cruzi.* To our knowledge, no modifications were found at D, except for acp3 in only one tRNA specie (Lys-TTT) from *T. brucei*. Thus, we hypothesize that the tRNA^Lys2(TTT)^ is also modified to acp^3^D47 in *T. cruzi*. The remaining 21 tRNAs, such as tRNA^Lys1A(CTT)^, tRNA^Asn1A(GTT)^ and tRNAVal1^(AAC)^, likely contain acp^3^U or acp^3^D at position 47.

*T. cruzi* has many paralogs of tRNA-modifying enzymes, which likely add complexity to the regulation of tRNA modification through several processes, including localizing modifications to distinct subcellular compartments, introducing modifications at distinct nucleoside positions, and differentially controlling modifications in the different parasite life forms. The presence of multiple modification enzyme paralogs in *T. cruzi* also highlights the intricate mechanisms regulating tRNA modifications and their potential impact on cellular processes. For example, the two paralogs of ABP140 (ABP140a and ABP140b) are responsible for 3-methylcytidine (m^3^C) in at least one of the following positions: C20, C32, or C47. In *T. brucei*, two paralogs, TRM140a (Tb927.10.1800) and TRM140b (Tb927.9.11750), have been identified as being associated with the m^3^C modification in tRNAs ^76,77^. However, only TRM140a is capable of modifying the m^3^C32 position ^77^, strongly suggesting that its homolog in *T. cruzi*, ABP140b, has a preference for m^3^C32 modification, while ABP140a may target other sites. ABP140b, but not ABP140a, showed decreased levels in the MT relative to the EPI form. According to the tRNA-seq datasets, RT-derived signals in C20 and C32 change in opposite directions. In the MT form relative to EPI and TCT forms, the signals at C20 increased whereas the signals at C32 decreased. Such distinct dynamic changes of tRNA modification may be attributable to the differential regulation of paralogs of tRNA modifying enzymes.

Our results showed that tRNA modification can control the differentiation frequency of *T. cruzi*. Suppressing the expression of Tyw1a enzyme and its associated modification OHyW promoted the transition from the non-infective EPI form to the infective MT form. One potential mechanism underlying this process is that the change in the nutrient availability triggers a reduction of OHyW37 on tRNA^Phe(GAA)^ could lead to the downregulation of certain proteins. During the metacyclogenesis, a reduction in global translation levels is crucial for EPI to MT differentiation ^78^. Additionally, the MT form translates a narrower range of proteins compared to the EPI form^7^ Therefore, the reduction of OHyW level potentially triggers a shift in the parasite’s gene expression required for the transition from the EPI to MT form.

One limitation of our study is the inability to distinguish between cytoplasmic and mitochondrial tRNAs. In *T. cruzi* both cytoplasmic and mitochondrial tRNAs are encoded in the nucleus and share identical sequences ^56,57^. Therefore, sequence data represents tRNAs from both compartments. This may explain why the frequency of some RT-derived signatures is relatively low. For example, tRNA^Trp4(CCA)^ undergoes C-to-U editing at position 34 only in mitochondria but not in the cytoplasm. We observed low-level misincorporation of U at this position, likely reflecting the low ratio of mitochondrial tRNA to cytoplasmic tRNA. Additionally, differential changes between cytosolic and mitochondrial tRNAs may mask each other. In *T. brucei*, position 37 in tRNA^Phe(GAA)^ is modified to OHyW in the cytoplasm and wyosine in mitochondria. Interestingly, the cytoplasmic Tyw1a is downregulated in the MT form, but mitochondrial Tyw1b is not changed, suggesting that this modification frequency is altered only in the cytoplasm. However, both OHyW and wyosine generate RT-derived signatures, and changes in one modification may be underestimated due to masking by the unchanged presence of the other. A modest change in the misincorporation signal at this position between EPI and MT forms could be attributed to this issue. Separating mitochondrial and cytoplasmic fractions during sample preparation will help resolve this issue.

Overall, we demonstrated that tRNA modifications are differentially regulated across the life stages of *T. cruzi* and identified specific tRNA-modifying enzymes linked to the parasite’s survival and differentiation. The data generated in this work lay the foundation for a deeper understanding of novel mechanisms of regulation of protein synthesis in trypanosomes that will be important for the development of new diagnostic tools and therapeutics.

## Supplementary Figures

**Figure S1.**
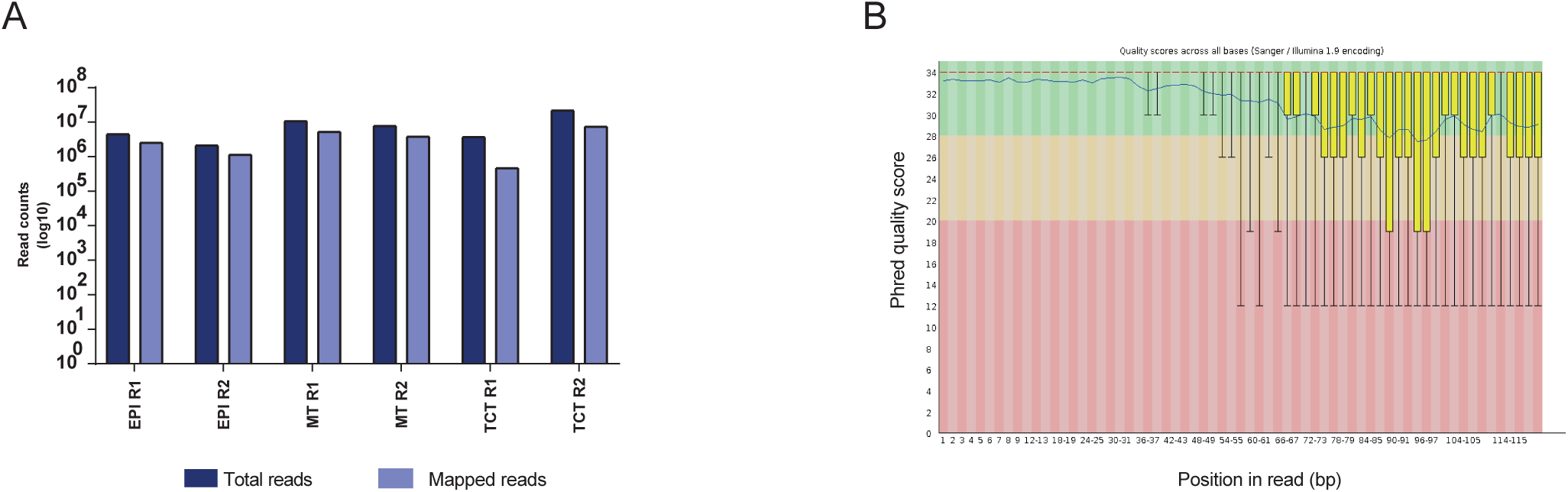
Quality check of tRNA-seq data in *T. cruzi* life forms (EPI, MT and TCT). **A)** Number of total reads and reads mapped to the *T. cruzi* genome are shown. Reads were exclusively mapped to tRNA sequences. **B)** Illustrative example of high sequence quality in the EPI R1 sample using Phred score as a parameter. R1= replicate 1 and R2= replicate 2.

**Figure S2.**
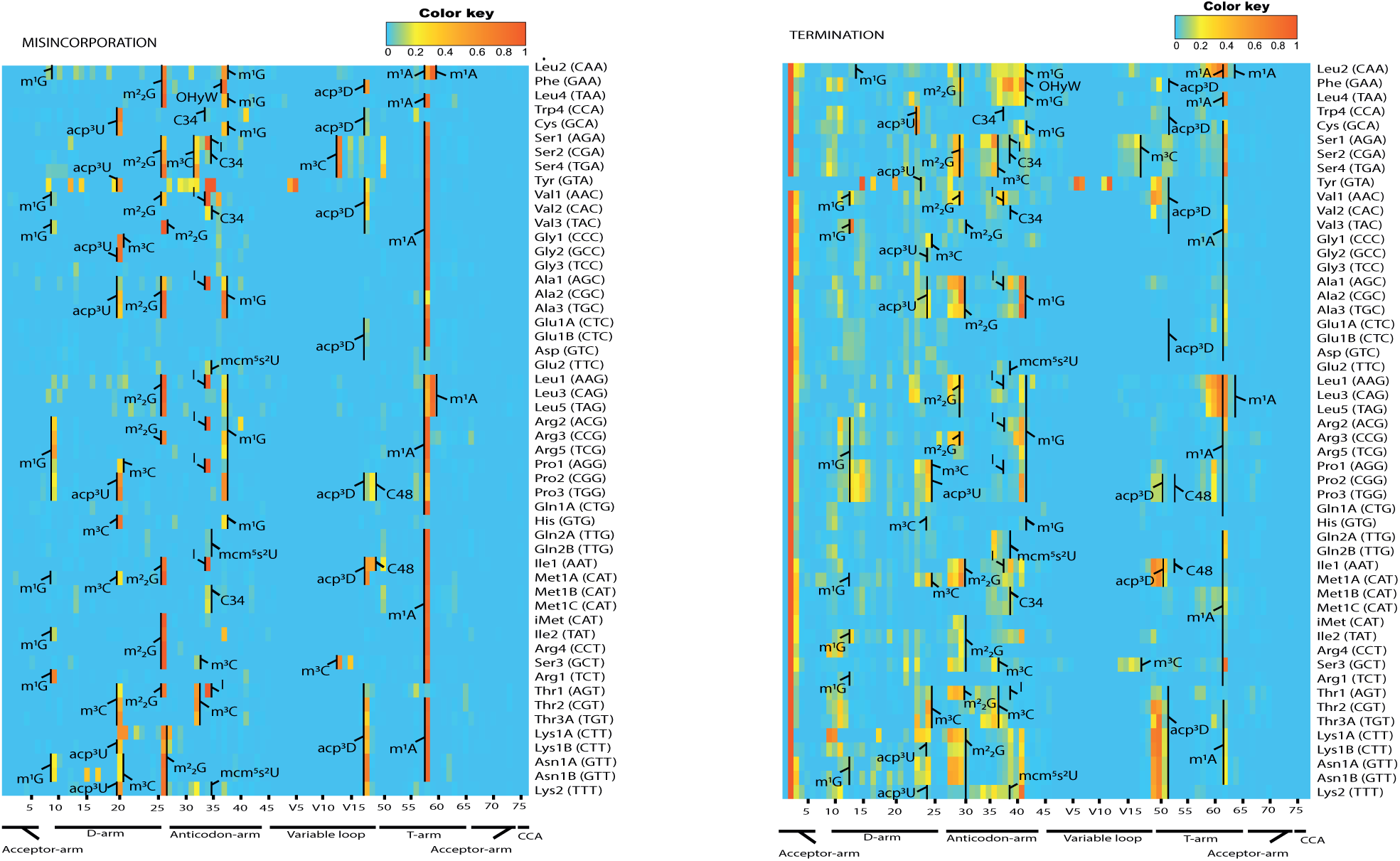
tRNA Modifications identified in EPI forms using tRNA-seq. The heatmaps showed the RT-derived signatures (Misincorporation and Termination frequencies) in the EPI R2 (second biological replicate) sample.

**Figure S3.**
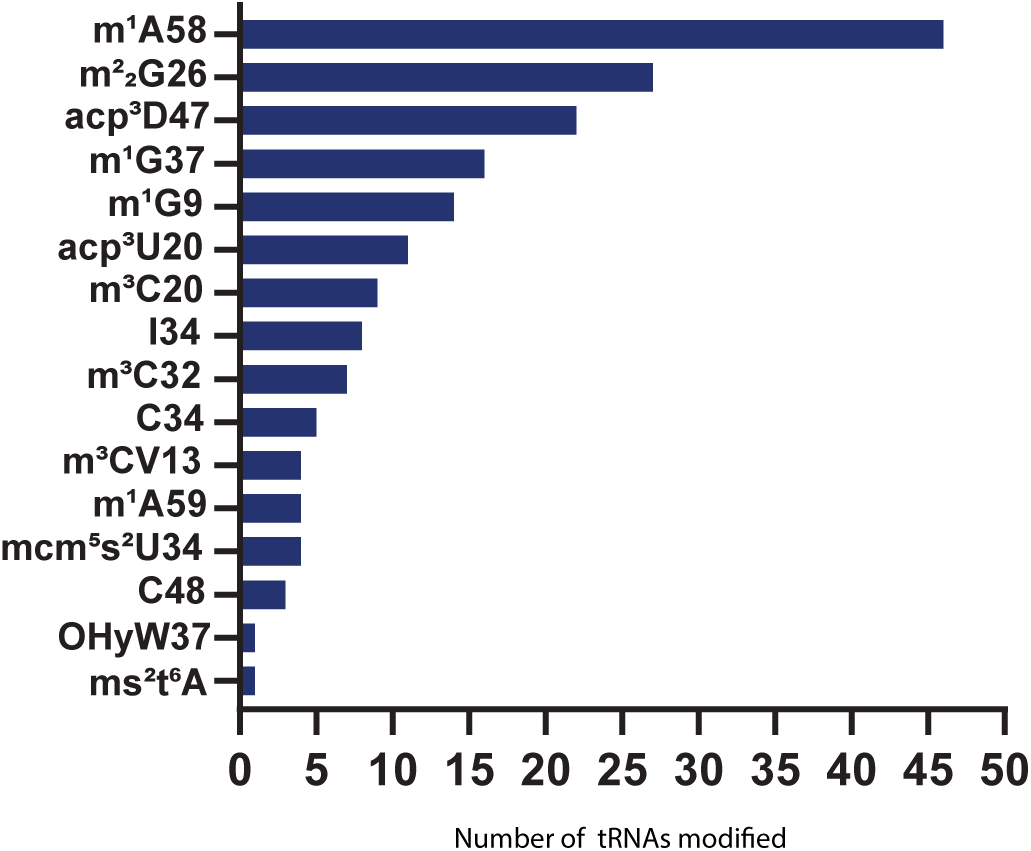
Number of tRNA modifications identified in different tRNA species from EPI using tRNA-seq approach.

**Figure S4.**
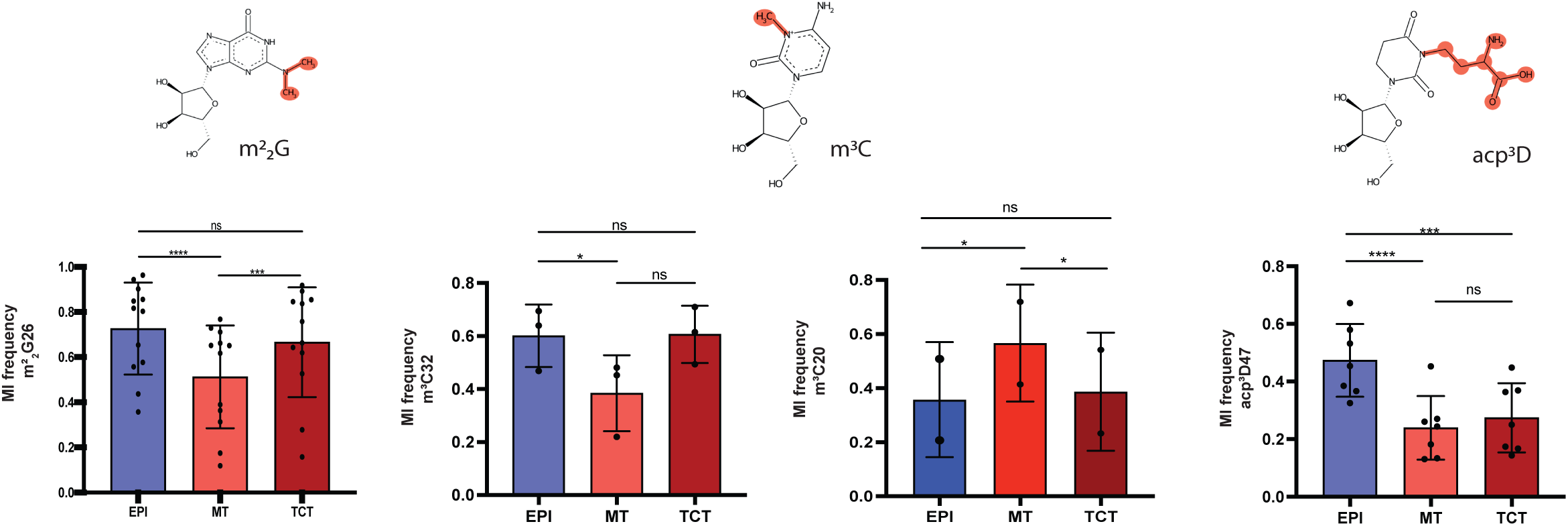
Misincorporation frequency of tRNA modifications that show differential abundance in at least one *T. cruzi* life form: m^2^_2_G26 (n=12), m^3^C32 (n=3), m^3^C20 (n=2) and acp^3^D47 (n=7). Statistical significance tests (p-value ≤ 0.05) were performed with the one-way ANOVA.

**Figure S5.** Generation and confirmation of Tyw1a knockout and add-back cell lines. **A)** Schematic shows the strategy used to replace the coding sequence of Tyw1a with blasticidin and puromycin resistance genes using the CRISPR-Cas9 technology as well as the oligonucleotides used for genotyping. **B)** Genotyping of five individual clones of the Tyw1a^-/-^ EPI cell line using the PCR strategies described in A. **C)** Graph shows relative Tyw1a mRNA levels of five individual clones of the TcTyw1a^-/-^ EPI cell line compared with the Cas9 parental cell line, assessed by RT-qPCR. **D)** Graphic representation of the vector used for generation of the Tyw1a addback cell lines. This vector allows the episomal expression of Tyw1a with a 6xHis tag in the C-terminal. **E)** Confirmation Tyw1a cloning in the 6xHis_pTEX_hygro vector using *BamH*I and *Hind*III restriction sites. **F)** Western blotting shows the expression of Tyw1a-6xHis (∼94 kDa) in three individual clones of the addback EPI cell lines. The loading control was the total proteins on the gel stained with trichloroethanol (TCE).

## Supplementary Tables

**Table S1**. Identification of tRNA-modifying enzymes in *T. cruzi* genome. **A)** List of the enzymes (gene ID) associated with tRNA Modifications in several eukaryotes and prokaryotes. **B)** BLAST results containing the best hit of the tRNA-modifying enzyme found in the *T. cruzi* genome, along with the associated tRNA Modifications. The proteins showing E-value ≤1e-10 were considered as tRNA modifying enzyme homologs. **C)** List of tRNA-modifying enzymes in *T. cruzi* and their orthologs in *T. brucei*. **D)** List containing the tRNA-modifying enzymes analyzed in public RIT-seq data ^44^ linked with *T. brucei* fitness during the differentiation of procyclic to bloodstream trypomastigote forms. **E)** Fold change and adjusted p-values for tRNA-modifying enzymes between EPI and MT forms obtained with Ribo-seq data.

**Table S2**. Characterization of tRNA Modifications in *T. cruzi*. **A)** Gene ID and genomic localization of all tRNA species present in the *T. cruzi* genome. **B)** List containing all tRNA Modifications either identified or predicted using tRNA-seq and BLAST, respectively. **C**) Values of proportion of misincorporation from tRNA-seq.

**Table S3.** Primer sequences and PCR conditions were used for the construction and validation of genetically modified *T. cruzi* parasites generated in this study.

## Data availability

The tRNA-seq can be accessed at the Sequence Read Archive (SRA) (https://www.ncbi.nlm.nih.gov/sra) under the following accession number: PRJNA1124437.

